# Identification of Target Objects from Gaze Behavior during a Virtual Navigation Task

**DOI:** 10.1101/2021.03.30.437718

**Authors:** Leah R. Enders, Robert J. Smith, Stephen M. Gordon, Anthony J. Ries, Jonathan Touryan

**Affiliations:** DCS Corp., Alexandria, VA, USA; DEVCOM Army Research Laboratory, Aberdeen Proving Ground, MD, USA; Warfighter Effectiveness Research Center; U.S. Air Force Academy, CO 80840

**Keywords:** Eye Tracking, Virtual Environment, Visual Search, Distractors, Dwell Time, Divided Attention

## Abstract

Eye tracking has been an essential tool within the vision science community for many years. However, the majority of studies involving eye-tracking technology employ a relatively passive approach through the use of static imagery, prescribed motion, or video stimuli. This is in contrast to our everyday interaction with the natural world where we navigate our environment while actively seeking and using task-relevant visual information. For this reason, vision researchers are beginning to use virtual environment platforms, which offer interactive, realistic visual environments while maintaining a substantial level of experimental control. Here, we recorded eye movement behavior while participants freely navigated through a complex virtual environment. Within this environment, participants completed a visual search task where they were asked to find and count occurrence of specific targets among numerous distractor items. We assigned each participant into one of four target groups: Humvees, motorcycles, aircraft, or furniture. Our results show a significant relationship between gaze behavior and target objects across subject groups. Specifically, we see an increased number of fixations and increase dwell time on target relative to distractor objects. In addition, we included a divided attention task to investigate how search patterns changed with the addition of a secondary task. With increased cognitive load, subjects slowed their speed, decreased gaze on objects, and increased the number of objects scanned in the environment. Overall, our results confirm previous findings from more controlled laboratory settings and demonstrate that complex virtual environments can be used for active visual search experimentation, maintaining a high level of precision in the quantification of gaze information and visual attention. This study contributes to our understanding of how individuals search for information in a naturalistic virtual environment. Likewise, our paradigm provides an intriguing look into the heterogeneity of individual behaviors when completing an un-timed visual search task while actively navigating.

## 1 Introduction

Active, unconstrained visual exploration is the sensory foundation of how the majority of individuals interact with the natural world, continually seeking information from their environment. This often includes coordinated body, head, and eye movement activity. In contrast, the majority of studies that seek to understand human visual perception employ a relatively passive approach through the presentation of stimuli, whether synthetic or natural. Likewise, body, head, and even eye movements are often constrained, either explicitly or by the nature of the experimental paradigm. These factors help control the manifold sources of variability, enabling the meaningful interpretation of finite empirical data. However, as both our understanding of perception and experimental capabilities expand, an increasing number of studies have sought to explore visual processes under more natural conditions; enhancing ecological validity while maintaining construct validity (Diaz et al., 2013; Foulsham and Kingstone, 2017). Here, we seek to extend previous work by quantifying the relationship between gaze behavior and target objects during active navigation of a complex virtual environment.

Measuring eye movement activity, including saccades, fixations, and blinks, has provided researchers a non-invasive way to gain valuable insight into perceptual, attentional, and cognitive processes during visual search tasks (Hoffman and Subramaniam, 1995; Kowler et al., 1995; Deubel and Schneider, 1996; Williams and Castelhano, 2019). Examining fixation patterns (e.g. number of fixations or dwell time) can indicate how individuals process visual information. For example, previous work has shown that individuals increase the number of fixations and dwell time (summation of all individual fixation durations) on informative visual objects within a scene (Loftus and Mackworth, 1978) and in the detection of changes of an object’s location within a scene (Võ et al., 2010). Improved memory recall and recognition on tasks is associated with increased number of fixations (Kafkas and Montaldi, 2011; Tatler and Tatler, 2013) and increased dwell time (Hollingworth and Henderson, 2002; Draschkow et al., 2014; Helbing et al., 2020). Specifically, increased number of fixations and increased dwell time on objects during visual search tasks are linked to improved memory for those objects (Hollingworth and Henderson, 2002; Tatler and Tatler, 2013; Draschkow et al., 2014; Helbing et al., 2020). This also appears to be the case when comparing how individuals visually attend to target objects compared to distractors in the environment. Horstmann et al. (2019) found that the average number of fixations on visual targets (about 1.55) was higher compared to the average number of fixations on similar looking distractors (about 1.20) during a search task with static images. Watson et al. (2019) reported that the number of fixations on targets ranged from about 3.3 to 4 compared to around 2.8 to 3.8 fixations on distractors, during a free visual search and reward learning task in a virtual environment. In terms of dwell time, Draschkow et al. (2014) found subjects looked about 0.6 seconds longer at targets as compared to distractors during visual search of static natural scenes.

Studying eye movement patterns can also provide information related to cognitive state, especially when assessed in conjunction with changes in pupil size (Hess and Polt, 1964; Van Orden et al., 2000; Chen and Epps, 2014). For instance, increased cognitive demand is associated with a decrease in blink rate (Benedetto et al., 2011; Maffei and Angrilli, 2018), a decrease in blink duration (Benedetto et al., 2011), and increased pupil dilation (Hess and Polt, 1964; Benedetto et al., 2011). Fatigue, in contrast, is associated with an increase in blink rate (Stern et al., 1994; Maffei and Angrilli, 2018), while saccade peak velocity is positively correlated with increased levels of arousal (Di Stasi et al., 2013). In addition, pupil dilation has been shown to track with the time course of decision making (De Gee et al., 2014; Cohen Hoffing et al., 2020). Thus, eye movement and pupillary measures are a rich source of information about cognitive processing and emotional state, and can be utilized to gain valuable insight into how individuals perceive, navigate, and interpret sensory information.

Previous work suggests that visual search tasks using traditional stimuli such as static pictures may yield different findings than those incorporating real world scenarios (Kingstone et al., 2003). Research has shown notable differences in gaze patterns between simple static versus complex dynamic visual search tasks, arguing for the increasing utilizing of dynamic scenes. For instance, Smith and Mital (2013) found increased dwell time on visual objects and increased saccade amplitude during a viewing and identification task in a dynamic scene compared to a static scene. We live in a visually complex world that includes many visual points of interest, depth, motion, and contextual scene information. Therefore, real-life environments are seemingly the optimal scenario to study naturalistic eye movement during visual search.

To this end, researchers have employed free navigation visual tasks in real-life scenarios such as walking outdoors (Foulsham et al., 2011; Davoudian and Raynham, 2012; Liao et al., 2019), walking indoors (Kothari et al., 2020), driving (Dukic et al., 2013; Grüner and Ansorge, 2017), and shopping in a grocery store (Gidlöf et al., 2013), to name a few. Although eye tracking in a real-life scenario allows free body movement, conducting studies in real environments can be difficult if not impossible to control; every subject’s unique actions makes a comparative analysis difficult. Fotios et al. (2015) noted this challenge in a study that examined movement patterns for pedestrians walking down the street. Examining eye movement patterns in real life environments also limits the design of the study in terms of the availability of targets and distractors (i.e. extant objects or limited by budget) and may be limited on the ability to gather neurophysiological measures such as electroencephalogram (EEG) recordings. Furthermore, real-world paradigms are often limited to only locally accessible environments and restrict researchers from studying more consequential scenarios where there are high demands for visual attention during a search task (e.g. looking for threat targets in a combat zone).

The use of virtual environments in perception research is an ecologically valid approach that provides the ability to conduct studies in an interactive but controlled dynamic environment (Parsons, 2015). Since eye-tracking systems can now be readily integrated with 3D rendering software (i.e. game engines), researchers can conduct eye movement studies in more realistic and immersive environments (Watson et al., 2019). Virtual environments also allow for research designs that may otherwise not be practical for a real-world implementation. For example, Karacan et al. (2010) utilized a 3D rendered virtual environment to examine shifts in gaze patterns as subjects repeatedly walked a loop path looking for isolated changes in the environment during each lap (e.g. a new object appearing, changing, and/or disappearing). The use of the virtual environment allowed for uninterrupted “physical” and visual inspection of an environment with tightly controlled visual changes. Virtual environments can accommodate research in attentional control and even allow for quantifiable interactions with objects in the scene. Helbing et al. (2020) utilized a virtual reality environment to examine memory encoding during target search of ten different complex and naturalistic indoor rooms. Furthermore, utilizing game engines as Unity3D (Unity Technologies), can allow for the participants to remain stationary during visual exploration of an environment and for researchers to perform synchronous acquisition of multiple physiological modalities, including respiration, electrocardiography (EKG), and EEG (Jangraw et al., 2014) that would otherwise be difficult in an ambulatory condition.

Researchers are beginning to take advantage of virtual environments to conduct navigation-based search tasks. Recently, Clay (2019) developed a virtual environment platform for the analysis of gaze behavior as subjects explored a large virtual enviornment. This environment included a variety of houses located within a intricate city street structure, and included a virtual sun to allow for natural lighting and give context for navigational cues. Subjects were instructed to observe the houses and later were ask to recall if they rememberered a subset of houses and their corresponding location. Researchers were then able to successfully show visual attention varied throughout navigation of the entire enviornment. Similarly, David et al. (2020) demonstrated the potential of eye tracking in virtual reality using visual search task with a gaze-contingent occlusion of either foveal or peripheral information. This group investigated how vision loss impacted head and eye movement when subjects were placed in a series of sixteen different virtual scenes and had 30 seconds to navigate and search through a room for a specific target object under one of three conditions: normal vision (no vision loss), simulated central vision loss, or simulated peripheral vision loss. They demonstrated similar findings to previous work in 2-D scenarios and found that head movement plays an important role in exploration of the environment. They also demonstrated the importance of head movement and peripheral vision (over central vision) during visual search. These works show the potential and efficacy of utilizing virtual reality and eye tracking to investigate visual perception under more naturalistic conditions using information-rich environments

Similar to these previous works, the current study seeks to isolate distinct gaze behaviors associated with target objects during an active visual search of a complex environment. Here, subjects freely navigate through a virtual world while completing a self-paced visual search task identifying assigned targets placed amongst many distractors (all other objects in the virtual environment other than targets). These distractors include a wide variety of objects that are, in some cases, similar in shape or color to the assigned target object. In addition, some of the participants are assigned a target category that includes more than one particular target object in the environment.

This study also includes an auditory divided attention task to increase participants’ cognitive load during a portion of the visual search task, which enables us to further investigate how participants compensate visual attention during a self-paced task in a complex environment.

The primary aim of this study is to demonstrate the feasibility of using more realistic stimuli (complex virtual environments) and tasks (simultaneous search and navigation) to investigate perceptual processes while maintaining a high level of experimental precision. To this end, we quantify the difference in gaze patterns between task-relevant targets and task-irrelevant distractors, during high and low cognitive load, comparing the results to previous studies which utilized more traditional visual search and encoding paradigms. Specifically, we expect there to be an increased number of fixations and dwell time on targets, as compared to distractors (Draschkow et al., 2014; Horstmann et al., 2019; Watson et al., 2019). We likewise expect subjects will visually explore targets at a closer distance as compared to other objects in the environment. Finally, we anticipate that auditory math task will elicit changes in saccade or fixation activity, such as increased visual attention on scanned objects in the environment (Pomplun et al., 2001; King, 2009; Buettner, 2013; Zagermann et al., 2018), or a change in general exploratory behavior (i.e. reduced speed or in the number of objects viewed) of the environment due to increased cognitive load during.

## 2 Materials and Methods

### 2.1 Subjects

Forty-Five subjects, recruited from the Los Angeles area, participated in this study (17 females with mean age ± standard deviation (SD) = 36.8 ± 12.3 years, 28 males with mean age ± SD = 41.6 ± 14.4 years). All subjects were at least 18 years of age or older and able to speak, read, and write English. All subjects signed an Institutional Review Board approved informed consent form prior to participation (ARL 19-122). All participants had normal hearing and normal or corrected-to-normal vision and had normal color vision. All subjects completed a web-based pre-screen questionnaire containing eligibility, demographic, and game-use questions. Additional color vision and visual acuity screening was conducted in-lab to ensure a minimum of 20/40 vision, using a standard Snellen Chart, and normal color vision, assessed with a 14-plate Ishihara color test. Any subject who did not pass the screening process was not included in the study. Subjects completed a simulator sickness screening questionnaire, the Simulator Sickness Questionnaire (SSQ) (Kennedy et al., 1993), before and after Pre-Test training (see below) and then again after the main study task. The mean and SD for the Total Weighted Score from the SSQ was 12.6 ± 16.4 before the Pre-Test, 32.1 ± 30.7 after the Pre-Test, and 38.1 ± 37.5 after the main study task. As part of the questionnaire, subjects answered questions relating to video game experiences and weekly usage of video games. The average number of years playing video games was 28.0 ± 11.4 years. The mean age when subjects began playing video games was 12.8 ± 7.1 years. Over half of subjects (51%) reported playing video games less than two hours a week. Almost a third (29%) of the subjects reported playing video games 2-7 hours a week. The remaining 20% of subjects reported playing video games for greater than 8 hours a week.

### 2.2 Procedure

During the experimental session, subjects participated in four separate tasks: a go/no-go serial visual presentation, an old/new recognition task, and two virtual environment tasks. However, only results from the virtual environment training and navigation tasks are described here. The stimuli in the other tasks were unrelated to the virtual environment.

### 2.2.1 Overview

Subjects were asked to freely navigate the virtual environment with the goal of searching for and counting their assigned target objects. All subjects were randomly assigned to one of four target categories: Humvee Group (N = 15 subjects), Motorcycle Group (N = 14 subjects), Aircraft Group (N = 9 subjects) or Furniture Group (N = 7 subjects). The Aircraft and Furniture Groups were introduced later in the data collection, which was eventually halted due to restrictions on in-person studies, hence the lower subject numbers. The aircraft and furniture targets were already present in the environment prior to introduction of the two new Groups, thus, all subjects in every Subject Group navigated the same environment with the same objects in the same order (Figure 1). Natural landscape features and trail markers provided a suggested path through the virtual environment (although subjects could freely explore in any chosen direction).

**Figure 1:**
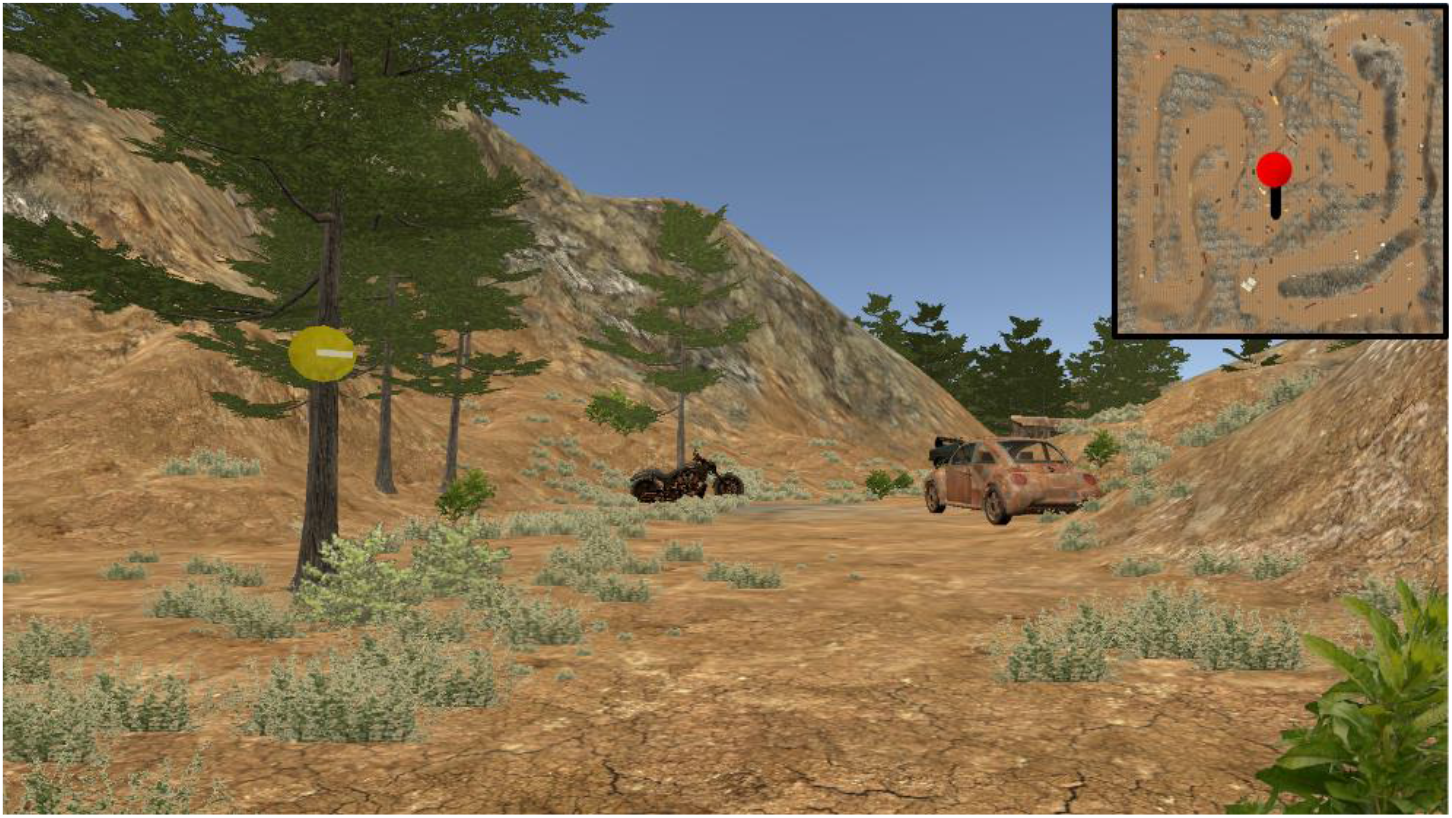
First person point of view near the beginning of the task. Trail makers (yellow circles with direction indicator) were placed on trees throughout the environment. Targets were assigned to each Subject Group to count: Humvee, motorcycle (shown), aircraft, and furniture. Distractors were any object in the environment not assigned to the subject (e.g. tires, dumpster, Humvees for anyone not in the Humvee Group). Inset (not visible during experiment) shows the current position on the complete map.

#### 2.2.2 System training task

A training task was used to acclimate subjects to navigation in the virtual environment via the keyboard and mouse. Movement was controlled with the W/A/S/D keys: “W” moved the subject in the forward direction, “A” allowed the subject to move left, “S” moved the subject backwards, and “D” allowed the subject to move right. A computer mouse was used to control the camera orientation or viewport (i.e. first person perspective) while in the virtual environment. This training environment was similar to the virtual environment used during the main task but contained different objects. This training task also ensured subjects were not acutely susceptible to simulator sickness.

#### 2.2.3 Testing setup

The experimental setup for this study combined multiple physiological modalities: eye-tracking, EEG, electrocardiography (EKG). Here, we described the relationship between task features, performance, and eye movement behavior. Other modalities, such as EEG and EKG, will be discussed in future reports and are not included in the current study.

All tasks were run using custom software built in the Unity 3D environment (Unity Technologies) run on the standard Tobii Pro Spectrum monitor (EIZO FlexScan EV2451) with a resolution of 1920 x 1080 pixels. Subjects were seated at a distance of approximately 70 cm from the monitor. Eye tracking data were collected with a Tobii Pro Spectrum (300 Hz). In addition to obtaining gaze position and pupil size, the Tobii Pro SDK was used to calculate the 3D gaze vector and identify the gaze vector collision object (in the Unity environment) for each valid sample. The eye tracking data were synchronized with the game state, keyboard, mouse, and EEG data using the Lab Streaming Layer protocol (Kothe, 2014). A standard 5-point calibration protocol was used to calibrate the eye tracker. The Tobii Pro Spectrum has an average binocular accuracy of 0.3°, binocular precision (root mean square) of 0.07°, and detects 98.8% of gazes (Tobii Pro, 2018). However, no verification of these error metrics was performed for this study. Head movement was not restricted in terms of head support or a chin rest. However, subjects were asked to maintain an upright, yet comfortable posture to minimize large upper body movements and maintain proper alignment with the eye tracker.

#### 2.2.4 Virtual environment description

Targets were placed in a pseudo-random sequence at semi-regular intervals along the path in the virtual environment. The location of all targets and objects in the environment were the same for all subjects. A general layout of the environment, indicating all target locations, is shown in Figure 2 and target examples are shown in Figure 3. Subjects all started at the same point on the virtual environment. Trail markers (N = 19) were placed along the trail for general navigational guidance. There were 15 targets total for each Subject Group. The same model of Humvee was used for all the *Humvee targets* and the same model of motorcycle was used for the *motorcycle targets*. For the *aircraft targets*, models varied and included helicopters, bi-planes, and one glider. For the *furniture targets*, targets included variations such as beds, grandfather clocks, tables, and a variety of seating furniture (e.g. sofa, dining chair). Sizes varied for the furniture with the chair being the smallest and the bed being the largest furniture target. Around 166 additional objects were included in the virtual environment that were not an assigned target to any Subject Group. These additional objects included, but were not limited to, cars, trucks, tanks, an oven, a drum set, a Ferris wheel, a pile of tires, dumpsters, and shipping containers. For analysis, a distractor was defined as any visual object in the environment not belonging to the specified Subject Group and included objects assigned as targets to other Subject Groups (e.g. Humvees were considered distractors for the Motorcycle Group). Terrain (e.g. trees, hillside, grass, path) and the sky were not included in the analysis unless explicitly mentioned.

**Figure 2:**
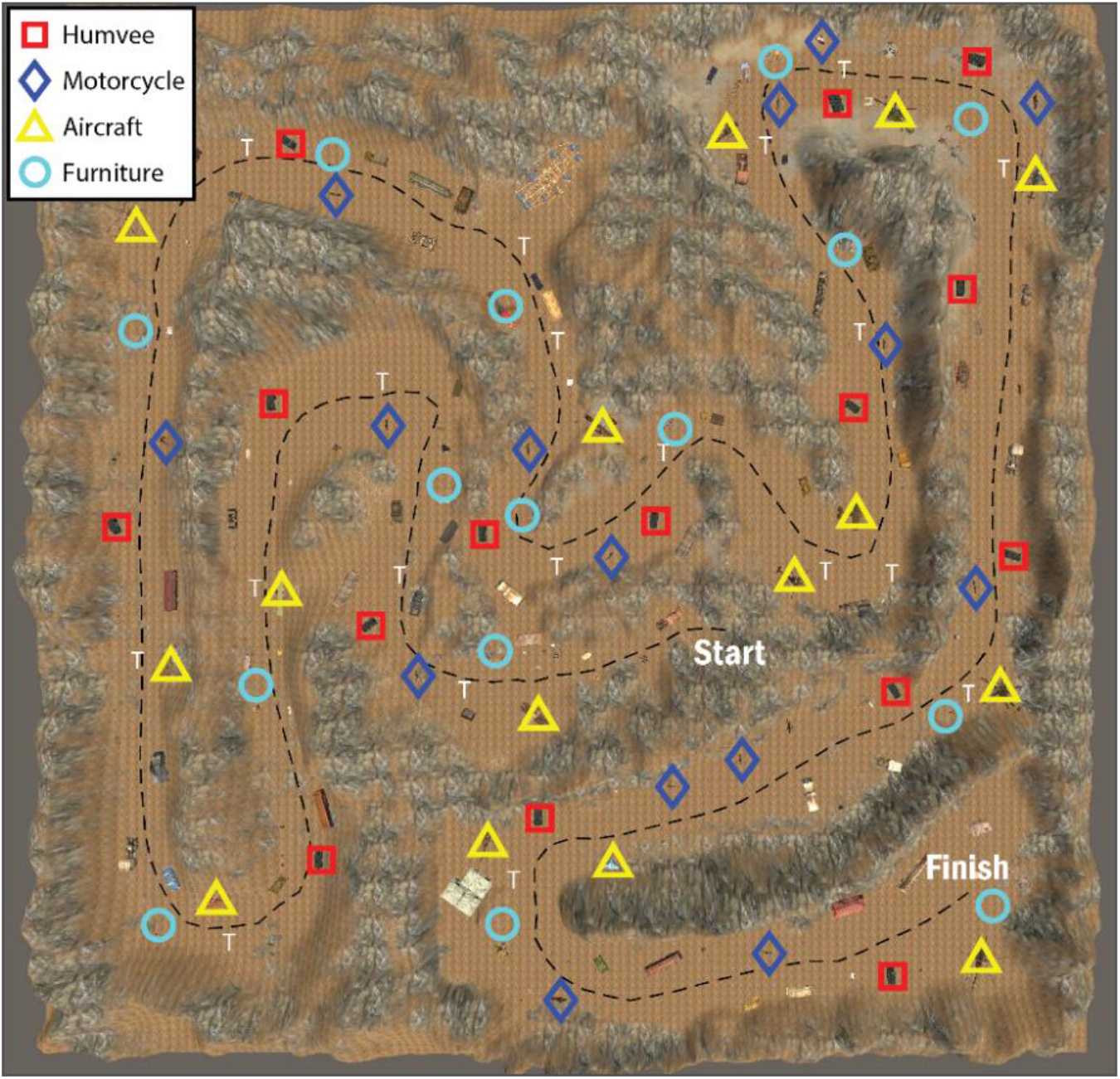
General layout of the virtual environment map. The black checkered line represents an example subject’s path from the starting area to the finish. Target icons are as follows: furniture (light blue circle), motorcycle (dark blue diamond), aircraft (yellow triangle), and Humvee (red square). Trail markers are present throughout the path and are indicated by a white “T”.

**Figure 3:**
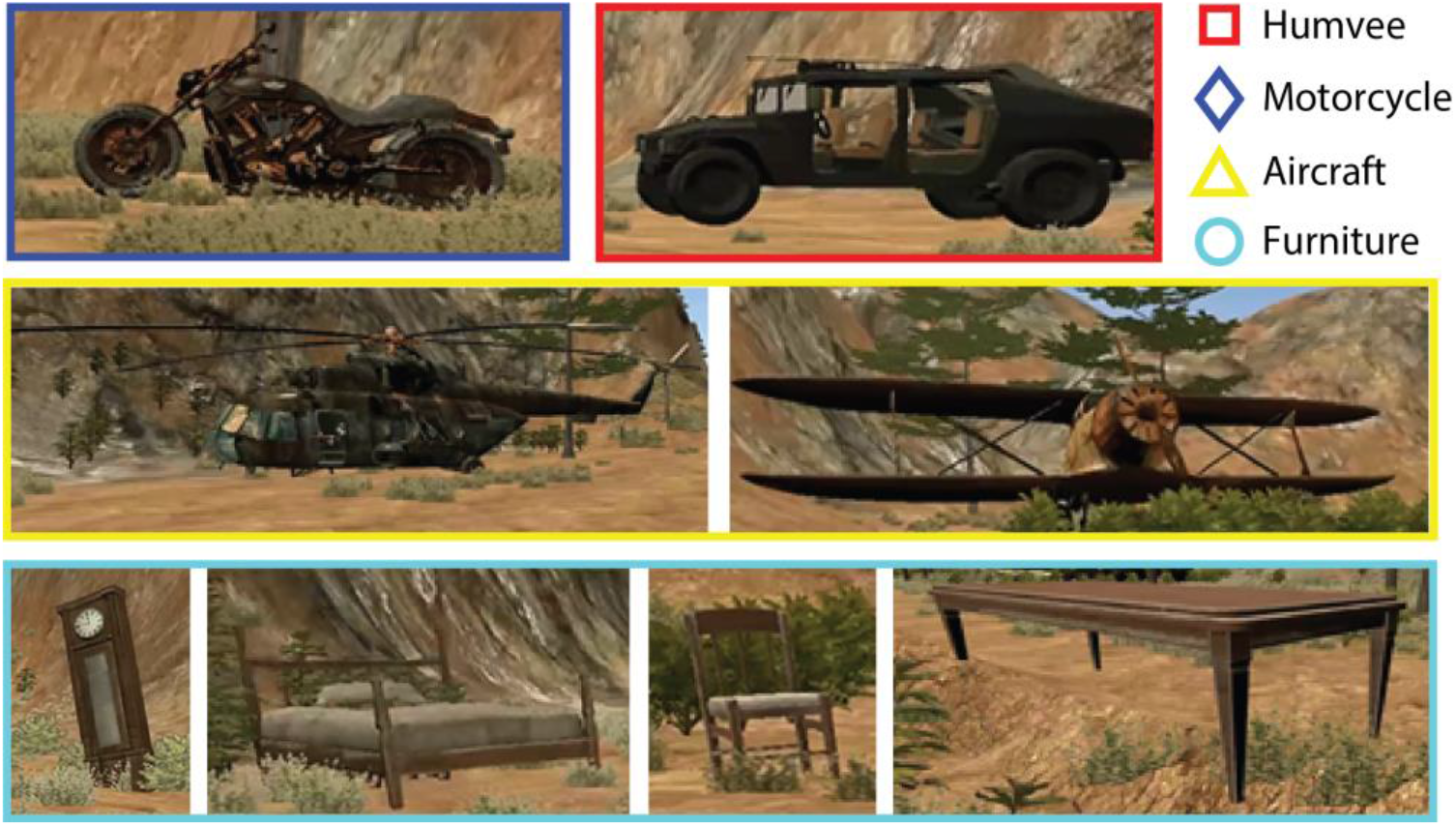
Example pictures of the targets for each Subject Group as they appear in the virtual environment. Motorcycles and Humvees (top row) did not vary in model but did vary in how they were positioned on the trail (i.e. motorcycle against a tree or at an angle by a rock). Both aircrafts (middle row) and furniture (third row) targets varied in shape, size, and positioning on the trail.

#### 2.2.5 Subject instruction and navigation

Subjects were instructed to search and count (mentally) when they saw a target assigned to their Subject Group. Subjects were encouraged to stay on or near the trail (and at times were verbally reminded by research staff) to make sure they encountered all objects, but were free to navigate as desired. Midway (8 min) into the session an auditory Math Task (divided attention task) was administered (see below for details). Subjects had up to 20 minutes to progress through the virtual environment and reach the finish. If subjects did not complete the task in 20 minutes, and if they did not encounter (as determined by their gaze vectors) at least 10 targets in the virtual environment, then their data was removed from statistical analysis. For this reason, data from two people in the Furniture Group were removed from all analysis. In addition, one subject in the Humvee Group reported feeling unwell during testing and experienced difficult navigating the environment (i.e. did not follow path) and thus, their data was also removed from the analysis. The average time to complete navigation of the virtual environment was about 12 (± 2) minutes. After completion, subjects were asked to recall how many targets they saw during the navigation task.

#### 2.2.6 Additional Math Task

Starting at the 8 minute mark, an auditory math problem was presented to the subjects. An auditory recording of a set of 3 to 4 numbers, with values between 0 and 9, was played for the subject through headphones (e.g. “4”, pause, “2”, pause, “8”, tone, subject reports “14”). A pause of 3 to 4 seconds separated each number in a set, and each set was followed by a tone. After the tone, subjects verbally reported the sum of numbers aloud to the experimenter. During the Math Task, subjects

were instructed to continue navigating through the virtual environment and continue searching and mentally counting their targets. This Math Task was repeated two more times (with different sets of numbers), for a total of three summation responses. There was an 8 to 30 second break between each set of numbers. Because the primary search task was self-paced, it is possible that a subject would finish exploring the virtual environment (reach the end of the path) without completing the Math Task. Only one subject (in the Humvee Group) did not complete the Math Task prior to finishing the navigation task and for this reason, their data was removed from the Math Task analysis.

### 2.3 Data extraction and analysis

#### 2.3.1 Fixation detection and object labeling

Blinks were identified from stereotyped gaps in the gaze position data (Holmqvist et al., 2011) while saccades (and corresponding fixations) were detected using a standard velocity-based algorithm (Engbert and Kliegl, 2003; Engbert and Mergenthaler, 2006; Dimigen et al., 2011). Specifically, we used a velocity factor of 6, a minimum saccade duration of 12ms, and a minimum fixation duration of 50ms, keeping only the largest saccade and subsequent fixation if two or more saccades were detected within the minimum fixation duration window. Visual inspection of five subjects’ first 500 saccades, shows the expected relationship between saccade peak velocity and saccade magnitude (i.e. main sequence) (Figure 4 a). These saccade distributions excluded blinks, dropouts, saccades with a duration shorter than 12ms and greater than 100ms, and peak velocities outside of a range of 25 and 1200 degrees per second. Visual inspection of the distribution of the saccade angle showed a strong tendency for subjects to scan the horizon (Figure 4b). Blinks were defined as fixation gaps with a duration ranging from 50ms to 500ms and dropouts were defined as any with a duration less than 50ms or greater than 500ms (any gap not considered a blink). After initial saccade detection, fixations of less than 100ms were discarded and not used in any subsequent analysis (Ouerhani et al., 2003; Mueller et al., 2008; Andersen et al., 2012). In addition to standard metrics associated with fixations (e.g. duration), each fixation was assigned a virtual environment object label using the following approach. Every valid gaze sample returned a corresponding object that was the result of the gaze vector collision. The object with the highest percentage of collisions over the fixation epoch was assigned as the “fixation object” (Figure 5 a). Target fixations were labeled as such if the highest percentage of collisions were on a target (e.g. motorcycles for the Motorcycle Group) and this number amounted to at least 10% of all gaze samples for that fixation. Distractor fixations were labeled using the same metric; receiving a distractor designation label if at least 10% of the gaze samples included the same distractor object. This 10% threshold was utilized to identify the primary fixation object and reduce the chance of including adjacent or background objects. Objects could be erroneously included in a fixation epoch from either gaze vector estimation error or having a relative position directly behind the primary fixation object. We selected the 10% value by assessing fixation labels at a range of thresholds: 0%, 1%, 10%, 20%, and 50% (Figure 5 b-c). The 10% threshold appeared to be a middle ground between reducing the chance of erroneous fixations without removing a large number of meaningful fixations. Fixation data from three subjects (two from the Aircraft Group and one from the Humvee Group) had a high dropout rate (high number of invalid samples). Thus these three subjects were removed from the analysis.

**Figure 4:**
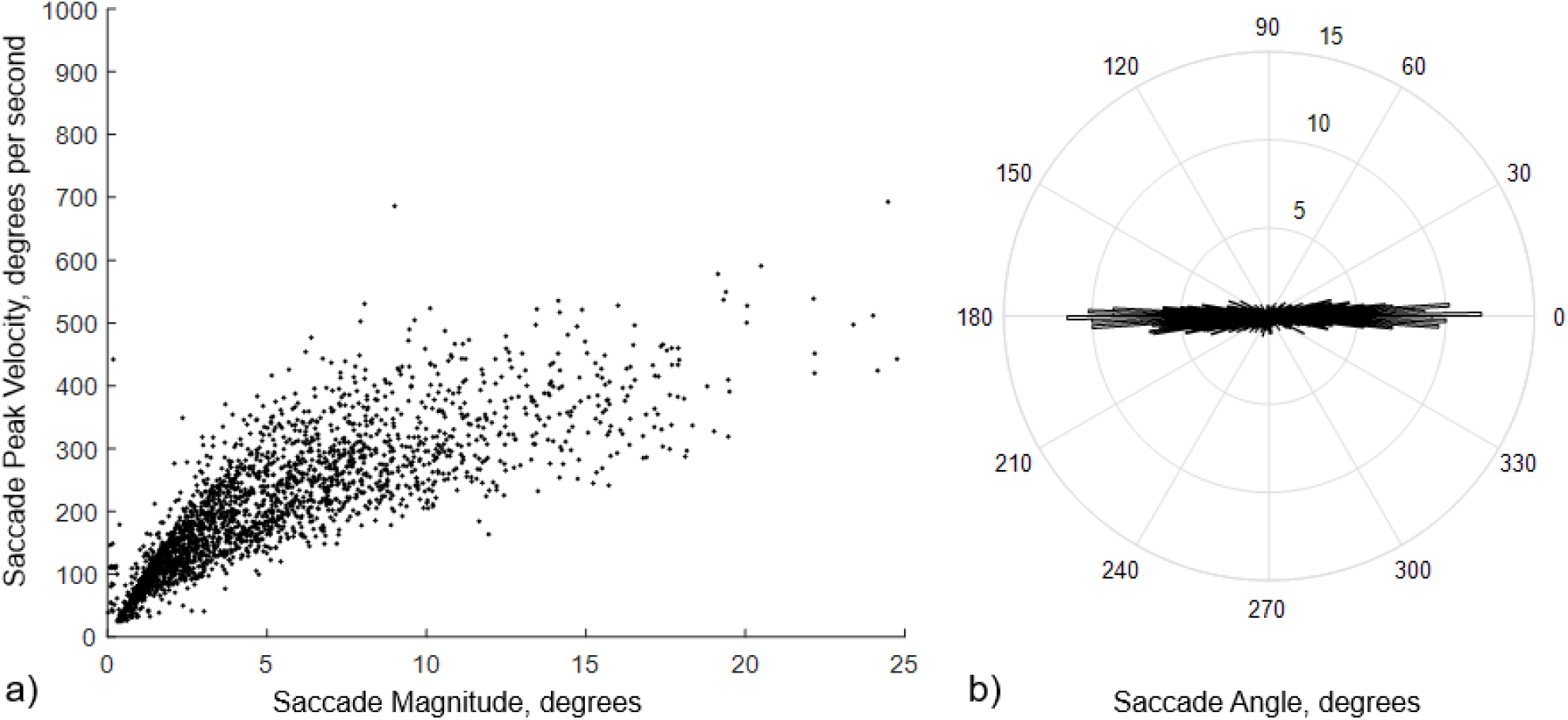
Saccade main sequence (a) and angle distribution (b) from a sample of the first 500 saccades from five representative subjects. The saccade angle distribution histogram has a one-degree resolution, with the radial axis showing average number and the angular axis showing average angle across all five subjects.

**Figure 5:**
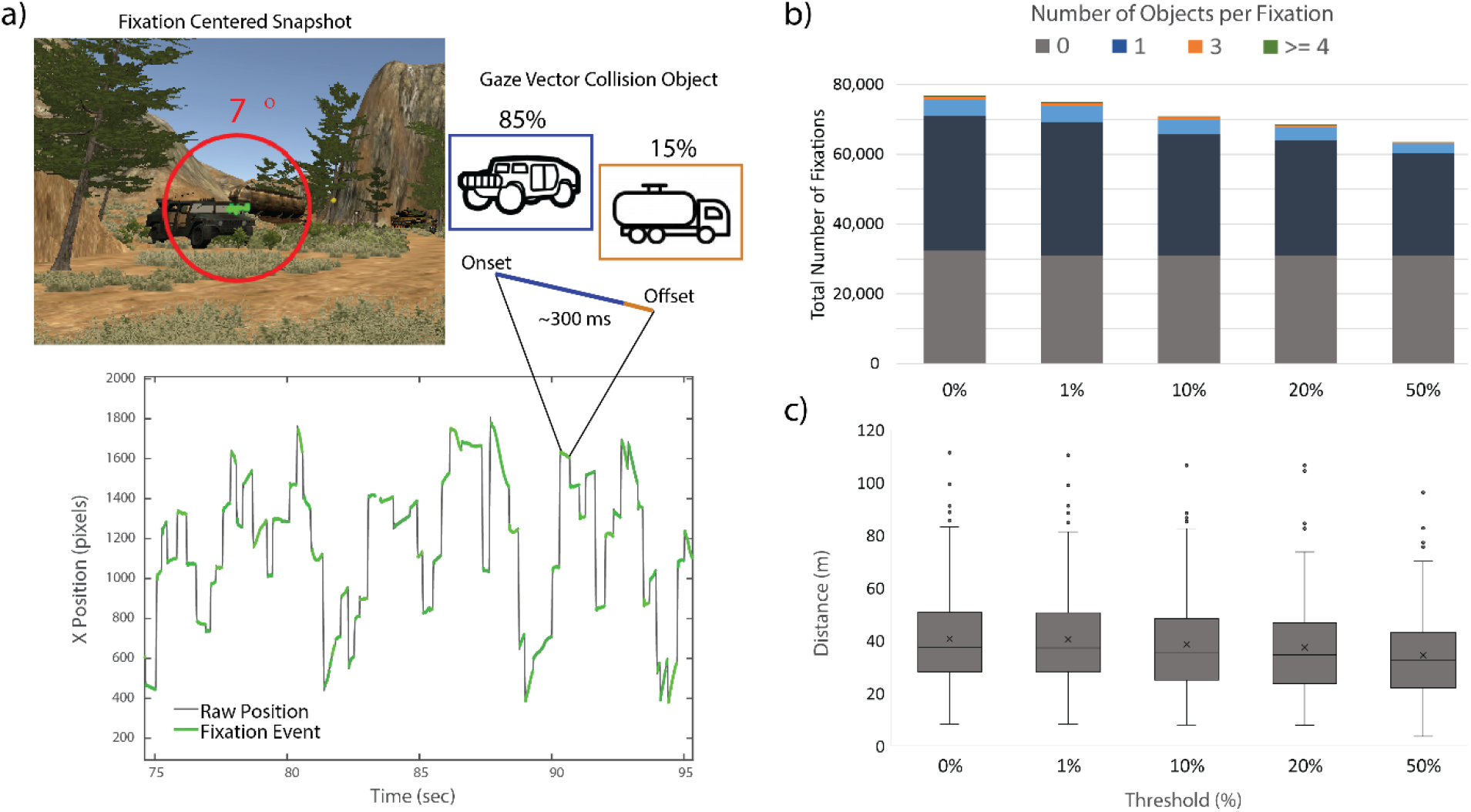
Fixation labeling approach. Saccades and fixations are detected from the raw X/Y gaze position time series. Each valid gaze sample is associated with an object via gaze vector collision. Fixations are then associated with objects if the dominant object (excluding terrain and sky) is contained in at least 10% of the samples of the epoch. (a) Alterative thresholds were examined and the 10% threshold appeared to filter out erroneous labels without excluding large amounts of meaningful data. The impact of the threshold value on the number of (c) and distance to (c) objects with all fixation epochs.

#### 2.3.2 Calculation of Fixation Variables for the main study analysis

From the fixation data for the targets, the following variables in Table 1, were calculated. The Self-Reported Target Count and the Gaze-Validated Target Count were calculated to compare subjective inventory with detected target fixations. To identify if our approach was sensitive enough to detect increased visual attention on targets, the Mean Number of Fixations, Mean Dwell Time, and Mean Distance were compared. Distance is included to provide a relative measure of how “close” subjects approached objects in the environment. Importantly, although the units here are given in meters, we acknowledge that this metric is not an equivalent analog to the real world (i.e. meters in the virtual environment may not reflect an actual meter in real life). Variation in object size and structural diversity impacted these particular fixation metrics. For instance, object surface area in the virtual environment was shown to be a large covariate with Mean Number of Fixations (Spearman’s rho =.719, *p*= .000), Mean Dwell Time (Spearman’s rho =.630, *p*= .000), and Mean Distance (Spearman’s rho =.558, *p*= .000). The larger the object, the increased chance a subject has to see it at any given viewing point, regardless of attentional focus. To help account for this bias, three additional variables, Normalized Number of Fixations, Normalized Dwell Time, and Normalized Distance were calculated utilizing the Global Number of Fixations, Global Dwell Time, and Global Distance variables. The global values were then used to normalize the means associated with the diversity of Target for each Subject Group.

**Table 1:** Dependent and independent variable list and definitions

| Variable | Definition |
| --- | --- |
| **Subject Group** | each subject was randomly assigned to one of four groups, named for the target they are assigned (i.e. Humvee Group assigned to look for Humvee targets) |
| **Self-Reported Target Count** | total count of targets that the subject reported seeing during exploration of the virtual environment |
| **Gaze-Validated Target Count** | total number of targets having at least one qualifying fixation associated with that target |
| **Mean Number of Fixations** | average number of qualifying fixations per each object |
| **Mean Dwell Time** | average total duration of fixations per each object |
| **Mean Distance** | average distance from the object when each associated fixation occurred |
| **Global Number of Fixations** | average number of fixations for that particular object across all subjects |
| **Global Dwell Time** | average dwell time for that particular object across all subjects |
| **Global Distance** | average distance from where that object was focused on across all subjects |
| **Normalized Number of Fixations** | Mean Number of Fixations subtracted from the Global Number of Fixations |
| **Normalized Dwell Time** | Mean Dwell Time subtracted from the Global Dwell Time |
| **Normalized Distance** | Mean Distance subtracted from the Global Distance |
| **Mean Duration of Individual Fixations** | average of all individual fixations across all Subject Groups and objects (targets and distractors) in the environment |
| **Fixation Rate** | summation of all fixations during the Math Task (or outside of the Math Task) divided by the total time spent in that time period |
| **Object Rate** | summation of the number of unique objects fixated on in the virtual environment divided by the total time spent in that time period |
| **Blink Rate** | summation of all blinks during the Math Task (or outside of the Math Task) divided by the total time spent in that time period |
| **Proportion of Fixations on Objects** | summation of fixations on objects (as opposed to terrain or sky) divided by sum of all fixations overall |
| **Position Velocity** | average change in position over time |

Due to uneven subject numbers across groups and the low subject numbers in the Furniture Group (see Limitations Section), the primary analysis in this report focused on the distinction between targets as compared to distractors, without attempting to compare across Subjects Groups. However, an additional analysis included a comparison of gaze data between just the Humvee and Motorcycle Groups to identify any difference in gaze behavior between the Humvee and motorcycle targets (referred to in the analysis as the independent variable, **Object Subset**). The Humvee and the Motorcycle Groups were utilized in this way because these two groups were comparable in subject numbers (N=13 and 14, respectively) and target attributes (i.e. same object model throughout the environment).

For the Math Task, fixation data from the following two time periods was compared: outside (before and after) and during the Math Task. To see if subjects compensated for divided attention during the Math Task by changing the rate at which they focus on objects, the Mean Duration of Individual Fixations *on objects* and Fixation Rate were compared between these time periods. To see if subjects compensated for divided attention during the Math Task by reducing the overall amount of visual attention devoted to each object, the Mean Number of Fixations per each object and Mean Dwell Time per each object in the virtual environment was compared between the two time periods. Object Rate was compared across the two time periods to capture visual scanning of different, unique objects in the environment. To see if subjects compensated for divided attention during the Math Task by reducing visual attention on particular objects and instead focused on background scenery, the Proportion of Fixations on Objects was compared between the two time periods. To see if subjects speed up or slowed down their progression of navigating through the environment, the Position Velocity was compared between the two time periods. Lastly, Blink Rate examined if subjects changed the number of blinks per unit of time with increased cognitive load.

### 2.4 Statistical analysis

To summarize, data from three subjects were removed due a high dropout rate, data from two subjects were removed due to not encountering the minimum threshold of targets, and data from one subject (who reported feeling ill during the testing) had navigational issues was removed, bringing the final inclusion of N = 39 subjects for analysis. In addition, one subject finished navigating the environment prior to the completion of the Math Task, for a total of N= 38 for that analysis. For the remaining subjects’, a normal distribution was assessed for all fixation variables using Kolmogorov-Smirnov and Shapiro-Wilk tests for normality. Parametric tests (i.e. Paired Samples t-test, MANOVA) were used for variables with normal distributions and non-parametric tests (i.e. Related-Samples Wilcoxon Signed Rank Test, Friedman Test) for non-normal distributions. For this reason, non-parametric statistical methods were utilized for the measures of Self-Reported Target Count, the Gaze-Validated Target Count, Blink Rate, and Position Velocity. All other variables had a normal distribution and parametric tests were used for comparative analysis. Outliers in the data were designated as samples/observations that were greater or less than three standard deviations from the mean. Outliers were removed from the data prior to analysis and includes one person’s data for Mean Distance and Normalized Dwell Time (N = 38 for analysis with these measures) and one person’s data for Mean Duration of Individual Fixations *on objects*, Mean Dwell Time *per object*, and Blink Rate during the Math Task analysis (N = 37 for analysis with these measures). In addition, Self-Reported Target Count was missing for six additional individuals (who did not report an answer when prompted) and one outlier was removed from the Self-Reported Target Count for a total of N =32 for analysis with this measure. A *p*-value of less than .05 was considered significant for all analyses and all analysis was conducted with IBM SPSS Statistics for Windows (Version 22, Armonk, NY: IBM Corp, Released 2013) software.

## 3 Results

### 3.1 Confirmation of fixated targets

On average, subjects reported the correct number of targets observed in the environment. A Related-Samples Wilcoxon Signed Rank Test compared the Self-Reported Target Count and the Gaze-Validated Target Count. There was no statistical difference between the two counts of the targets by subjects or identified by the system (Z = −0.573, *p* = 0.567). Median target counts were 15 for the Self-Reported and 14 for the Gaze-Validated.

### 3.2 General eye-gaze measurement outcomes

On average, individual fixations had a duration of about 0.32 seconds (320 ms) and the Fixation Rate was approximately 2.06 fixations-per-second throughout the main task when short fixations were removed. This is comparable to a Foulsham et al. (2011), who found an average of 2 fixations-per-second and an individual fixation duration of 441ms for subjects who watched a video of path they previously navigated. Subjects looked at objects (e.g. motorcycle, dumpster, trail markers) in the virtual environment, with a Mean Number of Fixations of 7.1 on each object. This is comparable to previous work by Zelinksy (2008) who found a Mean Number of Fixations of 4.8 on targets when searching for military tanks in a realistic scene. We found that subjects looked at objects with a Mean Dwell Time of 2.60 seconds per each object, which is comparable to work by Clay et al. (2019), who found a Mean Dwell Time of 5.53 seconds on visual objects for subjects who freely navigated and observed houses (considerably larger than most our targets) in a virtual environment town. We found that fixations on the surrounding terrain and sky compromised, on average, about 47% of all fixations. This is comparable Foulsham et al. (2011) who found that about 29% of fixations focused on the path ahead of where subjects were walking and Davoudian and Raynham (2012) who found about 50% of fixations were focused on the walking path.

Two separate one-way multivariate analysis of variances (MANOVAs) determined the effect of Fixation Object (target or distractor) on the normalized and non-normalized Mean Number of Fixations and Mean Dwell Time. There was a significant effect of Fixation Object for both non-normalized (F (2, 37) = 23.84, *p* = 0.000) and normalized gaze data (F (2, 36) = 22.54, *p* = 0.000) (Figure 6 a-b, c-d). Two separate Univariate analysis of variances (ANOVAs) examined how Mean Number of Fixations and Mean Dwell Time differed depending on Fixation Object. Subjects significantly increased both the Mean Number of Fixations (F (1, 38) = 35.73, *p* = 0.000) and the Mean Dwell Time (F(1, 38) = 48.84, *p* = 0.000) for targets compared to distractors (Figure 6 a&c). Two additional Univariate ANOVAs showed that Normalized Number of Fixations (F (1, 37) = 44.48, *p* = .000) and Normalized Dwell Time (F (1, 37) = 42.54, *p* = 0.000) also increased significantly for targets compared to distractors (Figure 6 b&d). Mean Distance was compared between Fixation Objects using a Univariate ANOVA (Figure 6 c). Subjects were significantly closer (less distance) to fixated targets compared to fixated distractors in the virtual environment (F(1, 37) = 12.99, *p* = 0.001). A separate Univariate ANOVA showed that Normalized Distance was also significantly less, on average, for targets.

**Figure 6:**
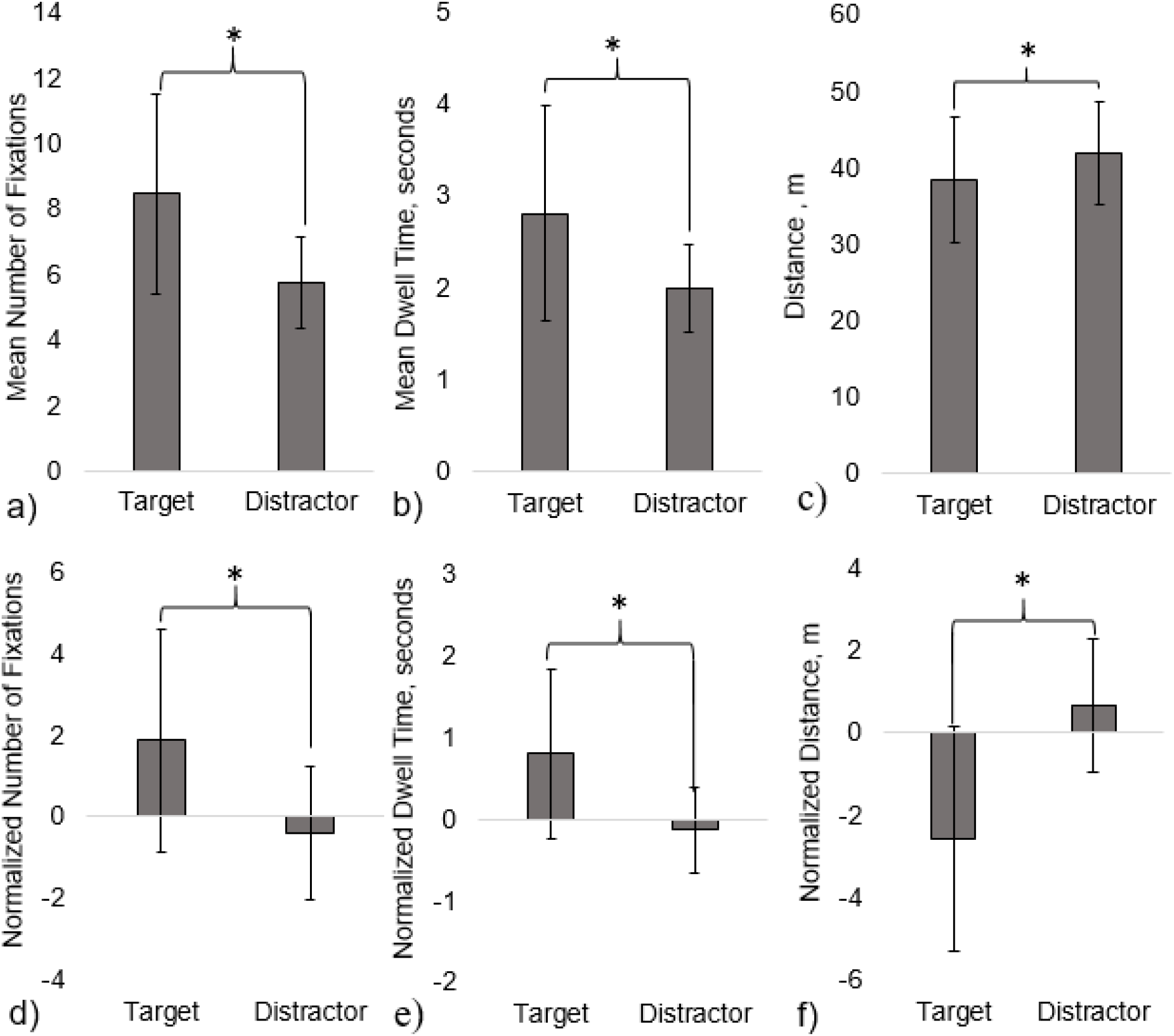
The Mean Number of Fixations (a), Mean Dwell Time (b), Mean Distance (c), Normalized Number of Fixations (d), Normalized Dwell Time (e) and Normalized Distance (f) were significantly greater for targets compared to distractors. Mean ± SD (error bars) are shown on graph.

### 3.3 Fixation differences for the Humvee Group and Motorcycle Group

Gaze data from the Humvee Group and the Motorcycle Group was used to quantify differences in the fixation patterns for the two target objects that were similar in terms of number, consistency, and dispersion along the path. Two separate two-way MANOVAs determined the effect of Subject Group and Object Subset (Humvee and motorcycles) and the interaction of Subject Group and Object Subset on the normalized and non-normalized Mean Number of Fixations and Mean Dwell Time. Overall, both non-normalized and normalized Mean Number of Fixations and Mean Dwell Time were significantly dependent upon the main effect of Object Subset and the interaction between Subject Group and Object Subset (Figure 7, Table 2). Four separate Univariate ANOVAs determined that the interaction between Subject Group and Object Subset was significantly different for non-normalized and normalized variables (Table 3). We found a significant main effect of Object Subset for the Mean Number of Fixations, where there was an overall greater number of fixations for the Humvees compared to motorcycles (Table 3). Object Subset was not a significant main effect for Mean Dwell Time, Normalized Number of Fixations, or Normalized Dwell Time. Tukey Post-hoc determined significant differences in those interactions. Both Subject Groups had significantly greater Mean Number of Fixations and greater Mean Dwell Time devoted to their targets, compared to the other object (*p*<.01, Tukey Post-hoc). Both Subject Groups had increased Mean Number of Fixations and Mean Dwell Time on their respective targets compared to that object for the other Subject Group (i.e. the Humvee Group focused on the Humvees significantly more than the Motorcycle Group focused on Humvees) (*p*<.01, Tukey Post-hoc). The same pattern of Post-hoc analysis statistical significance was found for the Normalized Mean Number of Fixations and Normalized Dwell Time (*p*<.05).

**Table 2:**
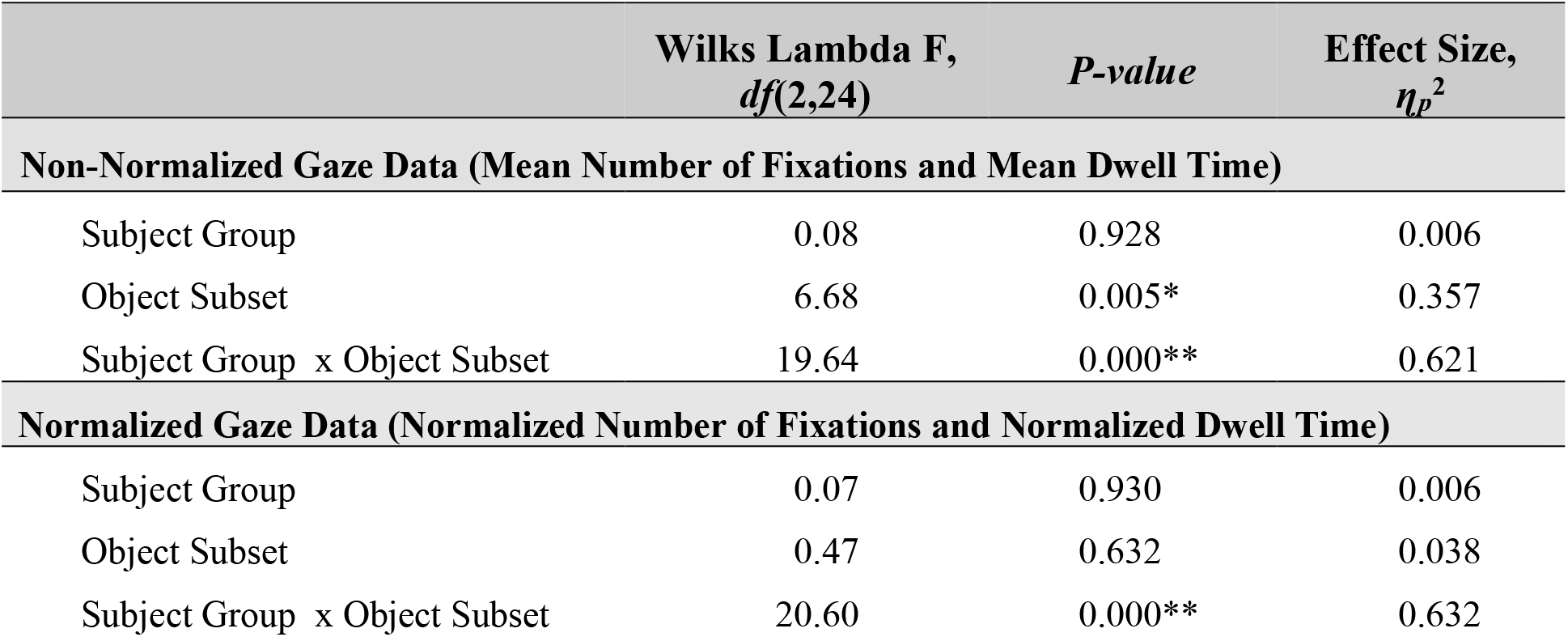
MANOVA for the non-normalized and normalized gaze data

**Table 3:** Univariate ANOVAs for the non-normalized and normalized gaze data

| | Wilks Lambda F,<br>*df*(1,26) | *P-value* | Effect Size,<br>$\eta_p^2$ |
| --- | --- | --- | --- |
| **Non-Normalized Gaze Data (Mean Number of Fixations and Mean Dwell Time)** |  |  |  |
| Mean Number of Fixations |  |  |  |
| Subject Group | 0.39 | 0.536 | 0.015 |
| Object Subset | 7.44 | 0.011* | 0.223 |
| Subject Group x Object Subset | 39.93 | 0.000** | 0.606 |
| Mean Dwell Time |  |  |  |
| Subject Group | 0.02 | 0.884 | 0.001 |
| Object Subset | 1.18 | 0.288 | 0.043 |
| Subject Group x Object Subset | 40.57 | 0.000** | 0.609 |
| **Normalized Gaze Data (Normalized Number of Fixations and Normalized Dwell Time)** |  |  |  |
| Mean Number of Fixations |  |  |  |
| Subject Group | 0.14 | 0.707 | 0.006 |
| Object Subset | 0.62 | 0.438 | 0.024 |
| Subject Group x Object Subset | 41.38 | 0.000** | 0.623 |
| Mean Dwell Time |  |  |  |
| Subject Group | 0.07 | 0.796 | 0.003 |
| Object Subset | 0.96 | 0.336 | 0.037 |
| Subject Group x Object Subset | 38.19 | 0.000** | 0.604 |

**Figure 7:**
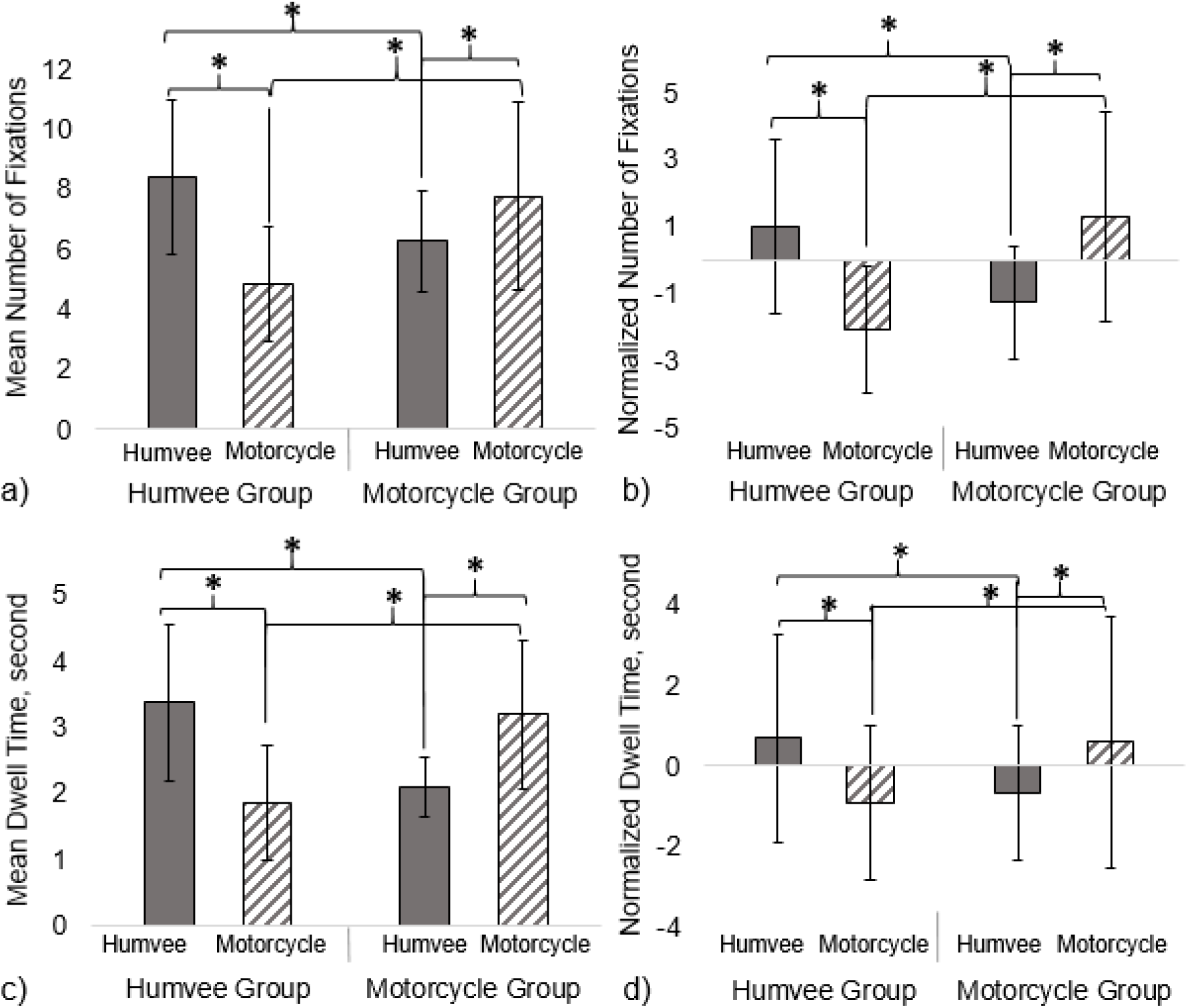
Both Subject Groups significantly increased Mean Number of Fixations (a), Normalized Mean Number of Fixations (b), Mean Dwell Time (c) and Normalized Dwell Time (d) for their respective targets. This increased Subject Group visual attention to targets was increased compared to that object for the opposite Subject Group. Mean ± SD (error bars) are shown on graph.

### 3.4 Effect of cognitive load

The total time spent on the Math Task was approximately 150 seconds (~2.5 min), compared to the time spent outside of the Math Task (before and after) 713 seconds (~12 min). Paired samples t-test determined that subjects did not significantly change the Mean Duration of Individual Fixations *on objects* (*t*_36_ = 0.03, *p*=.979) during the Math Task compared to outside of the Math Task. However, Paired samples t-tests showed that subjects significantly decreased their Fixation Rate (*t*_37_ = −2.91, *p* = 0.006) (Figure 8 c). This discrepancy was explained by the relative increase in Blink Rate during the Math Task (Related-Sample Wilcoxon Signed Rank Test, Z = 3.78, *p* = 0.000) (Figure 8 e). There was also a significant reduction in the Mean Number of Fixations *per object* (*t*_37_ = −5.67, *p* = 0.000) and the Mean Dwell Time *per object* (*t*_36_ = −4.51, *p* = 0.000) during the Math Task, as compared to outside the Math Task (Figure 8 a-b). In contrast, Object Rate increased significantly during the Math Task compared to outside the Math Task (*t*_37_ = 3.44, *p* = 0.001) (Figure 8 d). Interestingly, a Paired samples test showed that the Proportion of Fixations on Objects in the virtual environment (as opposed to fixations on terrain or sky) did not significantly change during the Math Task portion compared to outside of the Math Task (*t*_37_ = 0.16, *p* = 0.873). Additionally, a Related-Sample Wilcoxon Signed Rank Test showed that subjects significantly reduced their Position velocity, the speed at which they progressed through the environment, during the Math Task compared to outside the Math Task (Z = −4.87, *p* = 0.000) (Figure 8 f).

**Figure 8.**
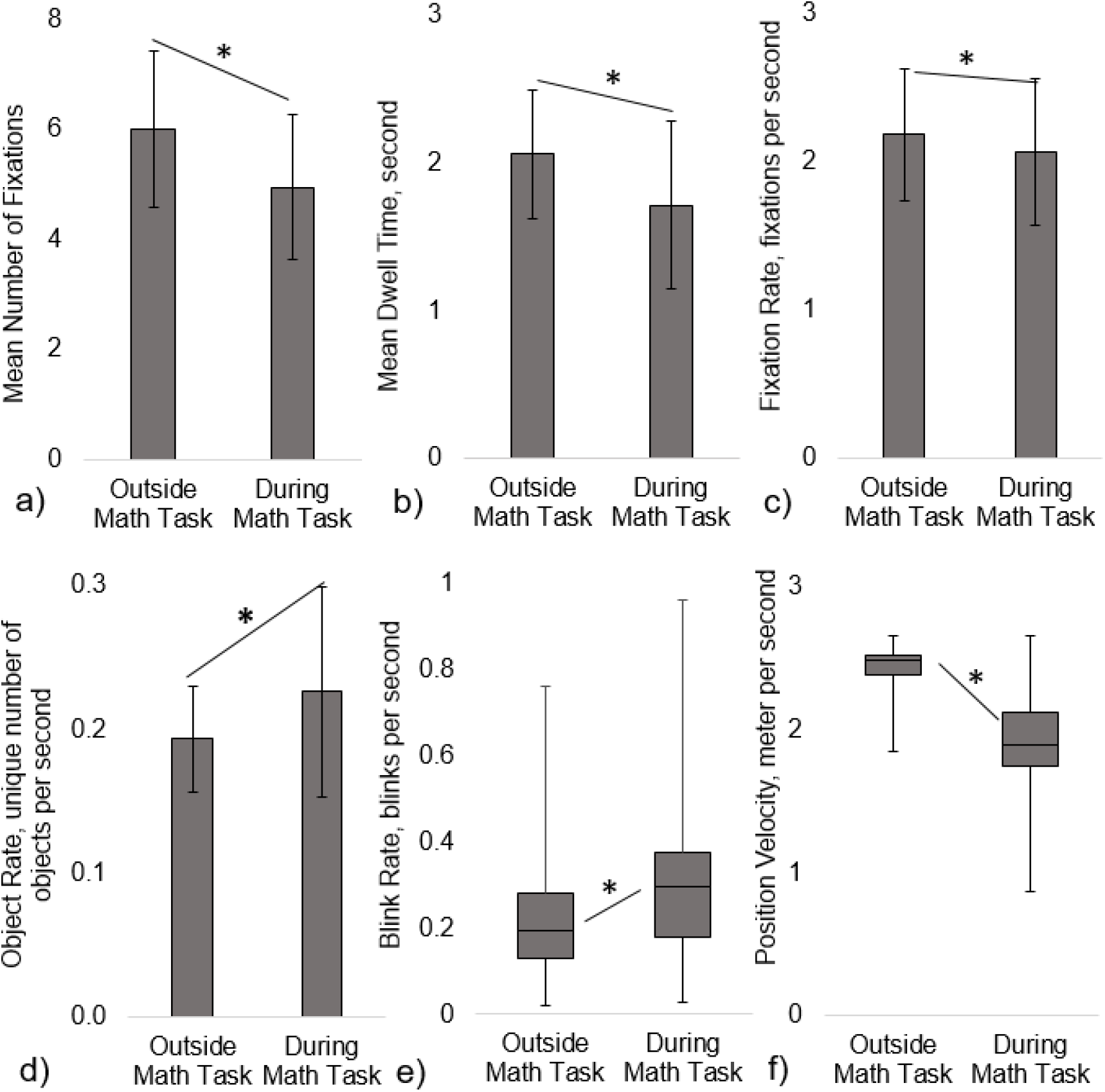
During the Math Task subjects significantly decreased the Mean Number of Fixations (a) Mean Dwell Time (b), and Fixation Rate on objects (c), but increased the number of unique objects fixated on per unit of time, Object Rate (d) and Blink Rate (e). Subjects significantly decreased their velocity navigating the environment (f) during the Math Task. The duration of individual fixations and the proportion of fixations on objects as opposed to terrain or sky, did not significantly change between time periods (not shown in figure). Mean ± Standard Deviation (error bars) shown on figures (a)-(d) and Median ± Interquartile Range (error bars) on figure (e)-(f).

## 4 Discussion

In this study, we demonstrate how a virtual environment can be used to identify subtle task-relevant gaze behavior in more a real-world context. Our approach enables us to collect meaningful, object-centered, gaze information during visual search in a complex environment without restricting virtual head movement (i.e. camera position and orientation). Consistent with previous studies, we show a clear distinction in gaze behavior between target and distractor objects. Moreover, we quantify how this gaze behavior changes when subjects’ attention is divided between visual search and secondary task.

### 4.1 General Discussion

Here, we observed a direct link between gaze activity and specific objects within the virtual environment. Overall, subjects looked at targets significantly more often and longer than distractors. This confirms our initial hypothesis, based on previous studies in more restricted experimental contexts, and demonstrates the feasibility of gaze analysis in dynamic (constantly changing) environments. This study is unique in that we acquired data from a relatively large number of subjects (N= 39) navigating a detailed and complex virtual environment, but were still able to identify distinct group-level gaze dynamics. This fixation and object-level precision would enable meaningful inferences from the concurrent use of neurophysiological signals such as EEG.

### 4.2 Increased Number of Fixations and Dwell Time on targets compared to distractors

In our study, we found subjects increased their visual attention (as measured by Mean Number of Fixations and Mean Dwell Time) on targets as compared to distractors. The Number of Fixations was, on average, 8.5 fixations for targets compared to 5.7 fixations for distractors. This 47% increase in fixations on targets over distractors was comparable to previous work. In a traditional visual search task with static images, Horstmann et al. (2019) found an approximate increase of 33% in Mean Number of Fixations for targets compared to target-similar distractors. Watson et al. (2019) found approximately a 10% increase overall in fixations for targets compared distractors in a study using a reward learning visual search task. We observed 41% increase in Mean Dwell Time for targets, with an average of 2.8 seconds, compared to Mean Dwell Time on distractors, 2.0 seconds. This outcome was comparable to work by Draschkow et al. (2014) who observed around a 30% increase in Mean Dwell Time on targets compared to distractors during a timed visual search task of complex static naturalistic scenes. Our result, showing increased overt visual attention on targets, supports our claim that such cognitive processes can be quantified in complex and dynamic virtual environments. Overall, our results were in line with previous studies, supporting the validity to our approach and processing methods. In addition, our unique virtual environments provides an intriguing look into the heterogeneity of individual behaviors when completing an un-timed visual search task while navigating.

It should be noted that the overall Mean Number of Fixations for both targets and distractors reported here, is greater than what has generally been found in many of the previous studies. This is likely due to the task design and nature of the environment. Even though subjects were given a maximum time of 20 minutes to complete the task, subjects were not instructed to find their targets as quickly as possible, as is the case in many visual search studies. Thus, subjects had more time to visually inspect all objects in the environment, without feeling rushed. Our environment contained 15 targets for each group and approximately 211 distractors (including other Subject Groups’ targets, a ratio of targets to distractors of about 1:14). Increasing the number of search items or the number of distractors can impact the working memory load and reduce visual search efficiency (Palmer, 1995; Wolfe, 2007, 2012; Zelinsky, 2008; Drew et al., 2017), especially in complex naturalistic environments (Wolfe, 1994b; Gidlöf et al., 2013). Therefore, the increased number of fixations observed in our study could be due to the subject’s self-pace progression through the environment and the particular target to distractor ratio.

Additionally, for some of our Subject Groups, target characteristics could have led to an overall high mean Number of Fixations on targets and distractors. For instance, distractors in some cases looked similar to the targets, especially at longer distances (i.e. Humvee vs another large vehicle). The effect of target-distractor similarity could have led to the need for increased visual attention to confidently distinguish between targets and distractors and decreased search efficiency (Duncan and Humphreys, 1989; Wolfe, 1994a, 2007; Zelinsky, 2008; Horstmann et al., 2019). It should also be noted that novelty of an object could have increased frequency of fixations. For instance, we would expect to see a difference in the Mean Number of Fixations and Mean Dwell Time for the Aircraft Group and the Furniture Group who had targets that varied in characteristics and models compared to the Humvee Group and Motorcycle Group with a target that stayed the same throughout the environment and only change in position and orientation in the environment. Subjects with a variable target may have fixated on more objects in general to determine if they should be included in their target count. Previous work has shown a disproportionate increase in visual attention on distractors for searches involving multiple targets compared single, static targets (Menneer et al., 2012). Novelty of the target can increase the time it takes to identify the object as a target among (varied) distractors (Lubow and Kaplan, 1997). The effect of target variation was not assessed in the current report due to low subject recruitment numbers in Subject Groups with a varied target. However, similar to previous work with multiple targets, we would expect that those with variable targets may have heightened attention towards distractors, negatively impacting their visual search efficiency throughout the task. Target characteristics, such as target variation and target-distractor similarity, may have been contributing factors to the large Number of Fixations reported overall.

### 4.3 Consideration of contributing factors due to task design

Inherent differences in visual objects’ shape, color, and size should have impacted visual attention towards specific objects in the virtual environment. However, rather than seeing these as limitations we argue that these are opportunities for additional, more nuanced research to better understand how: size, shape, color, visibility, context, etc. interplay with gaze behavior in ecologically valid environments. One would expect a greater Number of Fixations (and therefore greater Dwell Time) on the larger objects (e.g. Humvee, larger aircrafts, trucks, and buildings) compared to the smaller objects (e.g. furniture, motorcycles) due to being potentially visible at further distances. In contrast, a smaller object may be occluded by other larger objects or scenery until the subject is close to that object. In fact, we found that increased object surface size in the virtual environment was significantly and positively correlated with the Mean Number of Fixations, the Mean Dwell Time, and the Mean Distance from the object when the fixation occurred (see Materials and Methods). Therefore, it could be that participants naturally fixated more on the larger objects, even if such objects were not the target assigned to them and not relevant to their assigned task (Võ and Wolfe, 2012). This may have also given those assigned to Subject Groups with the larger targets, the Aircraft Group and the Humvee Group, a distinct advantage in seeing their targets due to visibility.

Along with size, visibility in terms of where the object was physically placed in the environment, may have also driven visual attention towards or away from some objects. Objects were sporadically placed throughout the environment and items placed at the end of long stretches of the path may have been central to subjects’ attentional locus while navigating down the path towards trail markers. These items, especially ones that were centrally located along the horizontal plane, may have naturally drawn more visual attention (Karacan et al., 2010; Foulsham et al., 2011), especially if they were a larger object. For example, we found a surprisingly high number of fixations (~16.5 fixations) and dwell time (~ 5.9 seconds) on a particular GMC truck located at the end of a long canyon before a tight turn (compared to 7.1 fixations and 2.6 seconds averaged for all objects). When examined further, this particular object also had the highest Mean Distance (~107 m) compared to the overall Mean Distance all objects in the environment (~ 40m). Therefore, some subjects could have fixated items due to their semi-random placement in the virtual environment rather than the due to the attributes of the item itself.

Scene context may also have impacted gaze towards certain objects in the virtual environment. For instance, the virtual environment was modeled as an arid and mountainous outdoor environment, but included some out of context items such as indoor furniture, musical instruments, a pool table, and a Ferris wheel. Scene context has shown to impact eye movement such as search time (Loftus and Mackworth, 1978; Henderson et al., 1999; Castelhano and Heaven, 2010) and memory recall (Draschkow et al., 2014). Items such as these may have garnered more visual attention due to their unexpected inclusion in the landscape (especially at the onset of the task) and/or could have been filtered as non-relevant visual objects if not assigned a Target Category that included those objects.

To help account for expected visual bias towards larger objects, random placement, or out of context objects in the virtual environment, we “normalized” the each fixation metric for every object by subtracting the global mean for that object (the averaged value across all participants for that particular object in the virtual environment). Normalization by simply dividing each gaze data point by the size of object (either 3D volume or 2D profile) in the virtual environment, resulted in a large bias towards the smaller targets. In contrast, our method of normalization enabled us to investigate object-centered gaze behavior for individuals compared to the mean across all groups for any particular object.

### 4.4 Discrepancy with virtual environment and real life walking scenario in distance of focus

Mean distance in the virtual environment was around 40 m, with fixations on targets occurring at closer distances than distractors. As noted previously, our virtual environment allowed subjects to view objects down the path or to look around to their surroundings. Here, subjects appeared to fixation on objects relatively further away in their environment, which was previously noted for studies measuring gaze in a virtual setting (Clay et al., 2019). However, we would expect there to be some discrepancy between our findings and what occurs in real-world ambulation. Foulsham et al. (2011) found that people focus on objects further away in the view field when watching a first person video walking through an environment, compared to when they walked that environment in real life. In an ambulatory scenario, gaze is more often focused on near-field objects or terrain that could potentially affect gait. In a virtual environment navigation, gait perturbation is not a factor, thus near-field obstacles may be “under viewed” compared to what would occur in the real world.

### 4.5 Effect of a divided attention task on gaze data

We found that subjects focused on more objects per unit of time during Math Task, not by changing fixation speed or shortening duration of each fixation, but by decreasing the Mean Number of Fixations on each object and therefore, total time spent on each object. Subjects also appeared to slow down their navigation speed (~24% decrease) and increase their Blink Rate (~46%) during the Math Task. Therefore, subjects appeared to compensate for increased cognitive load by reducing the number of times an object was scanned and slowing their pace of progression, rather than by shortening their fixation durations. Additionally, subjects did not appear to alter their visual attention away from objects and drift towards more background items in the environment (terrain and sky) when the auditory task was present.

This compensation method appears to be largely contrary to what others have found in studying eye movement and increased cognitive load during more traditional visual search experiments. Previous work has shown that as a task becomes more complex and difficult, there is an increase the Mean Number of Fixations (Pomplun et al., 2001; King, 2009; Buettner, 2013; Zagermann et al., 2018), an increase in Dwell Time (duration of fixations) (Pomplun et al., 2001; King, 2009; Meghanathan et al., 2015), an increase in the number of saccades (Zelinsky and Sheinberg, 1997; Zagermann et al., 2018) and an increase in saccade rate (Buettner, 2013). Importantly, we increased our cognitive workload by use of an auditory math task. There is a tendency to give attentional preference to auditory stimuli, potentially impacting one’s visual processing capabilities (Robinson and Sloutsky, 2010; Dunifon et al., 2016). In fact, previous neurophysiological work with EEG has shown that when auditory stimuli are paired with a visual task (cross-modal processing) there is a latency in the visual P300 response but no negative impact on the processing of auditory stimuli (Robinson et al., 2010). Pomplun et al. (2001) found reduced efficiency (increased reaction time, Number of Fixations, duration of individual fixations) in task performance when combining an auditory divided attention task with a visual matching task. Therefore, we would expect subjects to prioritize the Math Task and be less efficient in the visual search task. However, we observed almost the opposite effect of increased efficiency.

It should be noted the difference could be related to our tasking. First, our main target discrimination task was performed during active navigation (exploratory and self-paced) of an environment rather than a timed and speeded-response task. Since subjects could navigate the world freely without a tight time constraint, thus efficiency in visual search may not have been a priority, especially outside the Math Task. Second, our visual task was not difficult in terms of identification of their visual targets among the distractors (identify and add to overall mental target count) and only required subjects to continually keep a mental count. Third, we did not examine efficiency in terms of a difference in fixations on targets and distractors. Regardless, our findings provide additional insight into the effect of an additional auditory task during a self-paced visual search task in a natural virtual environment.

### 4.6 Limitations

We would like to recognize several potential limitations to our study. One limitation was a restriction in data collection efforts due to public health concerns; we had to cease data collection earlier than planned and so were unable to have a balanced number of subjects in each Subject Group. This resulted in limited capabilities for comparison among the Subjects Groups and their targets during the analysis. Second, while our experimental setup is similar to that of other studies, we utilized a desktop virtual environment instead of a virtual reality (VR) experience with a head mounted display. Although a VR system would provide a more immersive environment and allow for more free range in head and body movement compared to the current configuration, VR technology impose additional constraints when combining with other physiological measures, such as EEG. Likewise, simulator sickness is a common problem with immersive environments and our simulator sickness scores were relatively high overall. Simulator sickness could have impacted subject’s natural viewing process through an environment and act as an unintended distractor from the task. Finally, during the Math task it was observed that some participants paused navigation when listening to the auditory number presentation (~1-5 seconds), contrary to instruction and encouragement from experimenters. Therefore, gazed behavior during this time would be a reflection of cognitive processing and not necessarily the visual search and navigation task.

## 5 Conclusion

In conclusion, we found that even during a self-paced navigation of a complex virtual environment, eye movement data can be used to robustly identify task-relevant changes in gaze behavior. There was a significant relationship between a subject’s gaze behavior (Number of Fixations and Dwell Time), their target category, and objects in the environment. When an additional auditory Math Task was introduced, subjects slowed their speed, decreased the Number of Fixations and Dwell Time on objects in the environment, and increased the number of objects scanned in the environment. The present study adds to our understanding of how individuals actively search for information in a naturalistic environment.

## 6 Acknowledgement

We would like to thank Ashley Oiknine (AO), Bianca Dalangin (BD), Min Wei (MW), and Specifically, AO and BD for their work with subject recruitment and data collection, and MW for developing the initial world and task.

## 7 Funding

Research was sponsored by the Army Research Laboratory and was accomplished under Contract Number W911NF-10-D-0002. The views and conclusions contained in this document are those of the authors and should not be interpreted as representing the official policies, either expressed or implied, of the Army Research Laboratory or the U.S. Government. The U.S. Government is authorized to reproduce and distribute reprints for Government purposes notwithstanding any copyright notation herein.”

## Notes

### Competing Interest Statement

The authors have declared no competing interest.

